# Bridging the Dimensional Gap from Planar Spatial Transcriptomics to 3D Cell Atlases

**DOI:** 10.1101/2024.12.06.627127

**Authors:** Senlin Lin, Zhikang Wang, Yan Cui, Qi Zou, Chuangyi Han, Rui Yan, Zhidong Yang, Wei Zhang, Rui Gao, Jiangning Song, Michael Q Zhang, Hanchuan Peng, Jintai Yu, Jianfeng Feng, Yi Zhao, Zhiyuan Yuan

## Abstract

Spatial transcriptomics (ST) has revolutionized our understanding of tissue architecture, yet constructing comprehensive three-dimensional (3D) cell atlases remains challenging due to technical limitations and high cost. Current approaches typically capture only sparsely sampled two-dimensional sections, leaving substantial gaps that limit our understanding of continuous organ organization. Here, we present SpatialZ, a computational framework that bridges these gaps by generating virtual slices between experimentally measured sections, enabling the construction of dense 3D cell atlases from planar ST data. SpatialZ is designed to operate at single-cell resolution and function independently of gene coverage limitations inherent to specific spatial technologies. Comprehensive validation using real 3D ST and independent serial sectioning datasets demonstrates that SpatialZ accurately reconstructs virtual slices while preserving cell identities, gene expression patterns, and spatial relationships. Leveraging the BRAIN Initiative Cell Census Network data, we constructed a 3D hemisphere atlas comprising over 38 million cells, a scale not feasible experimentally. This dense atlas enables unprecedented capabilities, including *in silico* sectioning at arbitrary angles, explorations of gene expression across both 3D volumes and surfaces, and 3D mapping of query tissue sections. While currently validated for spatial transcriptomics, the underlying principles of SpatialZ could potentially be adapted for spatial proteomics, spatial metabolomics, and even spatial multi-omics. Validated through internal and external testing, our computationally generated atlas maintains biological accuracy, providing unprecedented resolution of spatial molecular landscapes and demonstrating the potential of computational approaches in advancing 3D ST.

## Introduction

Spatial transcriptomics (ST) has emerged as a transformative technology in modern biology, enabling researchers to map gene expression patterns while preserving crucial spatial information within tissues^1–3^. By capturing both transcriptomic data and spatial context, these technologies have revolutionized our understanding of tissue organization, cellular interactions, and gene regulatory networks in their native environment^4–7^.

Recent advances in ST have primarily focused on two-dimensional (2D) tissue analysis^8–12^, where individual planar tissue sections are processed to generate cell atlases with transcriptomic and 2D spatial information^4,13–17^. While several laboratories have attempted to extend these capabilities to three-dimensional (3D) analysis using thick tissue blocks^18–20^, current approaches face significant limitations. Existing 3D ST technologies are restricted to relatively thin tissue sections (up to 200 micrometers in most recent advances^18^), making organ level 3D mapping impractical. The complex experimental protocols and resource-intensive nature of these approaches further limit their adoption in the wider research community.

To address these challenges, computational methods have been developed to align serial 2D sections within a common 3D coordinate system^21–24^. This approach has enabled the integration of extensive 2D ST datasets, including diverse organ cell atlases, cancer studies, and brain cell atlases generated by the BRAIN Initiative Cell Census Network (BICCN)^25^. However, these methods represent an initial step towards 3D, since they essentially create a stack of 2D sections rather than a 3D volume, leaving substantial gaps (e.g., 100 micrometers) between adjacent sections unmeasured. These gaps create discontinuities in the spatial information and prevent comprehensive 3D analysis of gene expression patterns and organ-level architecture.

An ideal solution for bridging the gap between 2D and 3D spatial transcriptomics should address several key requirements: (1) generation of comprehensive 3D organ atlases covering substantial portions of the organ, (2) independence from gene coverage limitations of specific ST technologies, (3) single-cell resolution capability, (4) accurate representation of 3D gene expression patterns on both surface and volume, (5) identification of 3D tissue regions and spatial domains of interest, and (6) ability to perform *in silico* sectioning for generating additional 2D slices at any desired angle or thickness.

In this study, we introduce SpatialZ, a computational framework that constructs a “pseudo 3D atlas” from sparsely sampled 2D tissue sections. Drawing inspiration from “pseudo-time” in single-cell analysis^26^—where time is inferred from gene expression continuity rather than actual temporal measurements—our “pseudo 3D atlas” reconstructs three-dimensional tissue architecture based on the continuity of niche transition across sections. While current technological limitations prevent the generation of real experimental 3D atlases, SpatialZ approaches this gap by interpolating between measured 2D sections to generate “virtual slices,” ultimately creating a dense and 3D spatial cell atlas.

We conducted comprehensive evaluations to validate SpatialZ’s performance. Using a real 3D spatial transcriptomics dataset, we simulated serial sectioning and demonstrated SpatialZ’s ability to accurately recover held-out sections, validating through gene expression patterns and spatial organization. We further confirmed the reliability of our approach using an independent 2D serial sectioning dataset, which revealed strong consistency between real and virtual slices and demonstrated enhanced spatial domain identification after virtual slice integration. We then applied SpatialZ to construct a pseudo 3D atlas of the mouse brain using a recently generated spatial atlas containing 129 sparsely sampled sections. Thanks to SpatialZ’s computationally efficient design, we constructed this atlas of 38,044,533 cells in 801 hours, which is not feasible experimentally. The resulting pseudo 3D atlas enabled three key capabilities previously impossible to achieve experimentally or computationally: *in silico* sectioning at arbitrary angles and positions, direct visualization of gene expression on both 3D volume and surfaces, and searching and locating a query tissue section within the 3D spatial atlas. We validated these computationally generated slices through comparison with in situ hybridization images, confirming the biological accuracy of our approach. Moreover, the pseudo 3D atlas generated 10 times more cells than the original experiment, complete with cell type and spatial domain annotations, providing comprehensive molecular and spatial organ architectures. While currently validated for spatial transcriptomics, the underlying principles of SpatialZ could potentially be adapted for spatial proteomics^27–31^, spatial metabolomics^32–35^, and even spatial multi-omics^36–40^. SpatialZ represents a powerful tool for understanding complex tissue architecture and gene expression patterns in three dimensions.

## Results

### SpatialZ overview

SpatialZ is a computational framework to construct a “pseudo 3D atlas” from sparsely sampled 2D tissue sections. Due to technical limitations, current spatial transcriptomics technologies face significant challenges in generating real three-dimensional organ cell atlases. While serial sectioning approaches provide valuable spatial information, the substantial gaps between consecutive sections (e.g., 100 micrometers) create discontinuities that hinder comprehensive 3D analysis. Drawing inspiration from pseudo-time analysis in single-cell analysis, where temporal trajectories are inferred from the continuity along the gene expression manifold^26,41,42^, SpatialZ reconstructs three-dimensional tissue architecture by leveraging continuity of niche transition between adjacent sections. The framework processes gene expression profiles and spatial coordinates from paired tissue sections, integrating cell type information and other annotations (e.g., spatial domain, if available) to generate intermediate “virtual slices” that bridge the gaps in z-axis sampling (Fig. 1a and Methods).

**Fig. 1.**
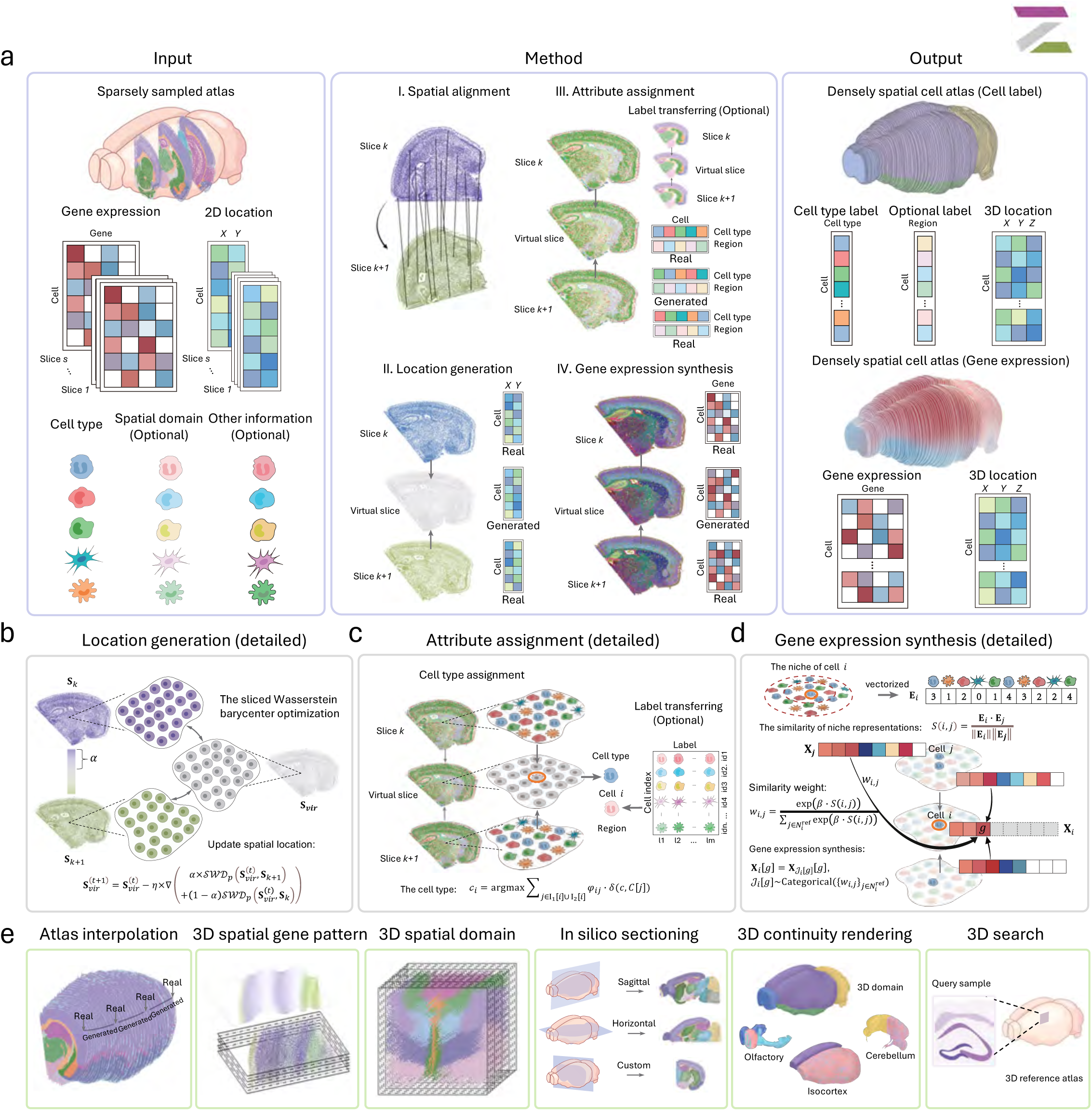
Overview of the SpatialZ workflow. **a,** SpatialZ takes the sparsely sampled atlas comprising the spatially resolved gene expression and location matrixes, cell type annotations, and other attribution (e.g., spatial domain information, if available) as inputs (left). (I) Spatial alignment of slices under a common coordinate system. Three key functional modules of SpatialZ for spatial brain atlas generation, including (II) location generation; (III) attribute assignment and (IV) gene expression synthesis (middle). SpatialZ interpolates the generated virtual slices between measured 2D sections, creating a dense spatial cell atlas including 3D location, restored gene expression matrixes and attribute information (right). **b-d**, Technical details of location generation (**b**), attribute assignment (**c**), and gene expression synthesis (**d**). **e**, Schematic of SpatialZ applications, including sparely sampled atlas interpolation, 3D spatial gene pattern identification, 3D spatial domain clustering, *in silico* sectioning, 3D continuity rendering for gene expression in specific field of view (FOV) tissue and 3D spatial search.

The SpatialZ workflow comprises four integrated computational steps. First, adjacent tissue sections are aligned within a common coordinate system using established spatial registration methods such as PASTE^24^ or reference frameworks like the Allen Mouse Brain Common Coordinate Framework version 3 (CCFv3)^43^. This step is done as traditional approaches of building a stack of 2D slice atlases, to establish the skeleton for 3D reconstruction. Second, to generate cell locations on virtual slices, SpatialZ employs sliced Wasserstein barycenter optimization to determine the number of cells and the spatial localization of all the cells in virtual slices. The number of interpolated slices can be either automatically calculated based on physical spacing or manually defined by users (Fig. 1a, 1b and Methods). This step ensures that the cell spatial localization distribution of the middle virtual slice represents a transitional state between the cell spatial localization distributions of the upper and lower sections. Third, to establish the cell type labels in each virtual slice, SpatialZ designed a sampling strategy based on the cell type composition within two nearest cell niches from the adjacent real sections (Fig. 1a, 1c). This step ensures that the cellular composition of each niche in the middle virtual slice shows a continuous transition of cell type proportions of the nearest niches from the upper and lower real sections. Our hypothesis is that it is cell niches that are continuous along the physical z-axis, not individual cell types. Finally, gene expression profiles for virtual cells are synthesized using a weighted sampling approach based on niche similarity with neighboring cells of matching cell type (Fig. 1a, 1d). This step ensures that the generated gene expression profiles simultaneously consider both the cell niche context and cell type identity, while avoiding the generation of artificial gene expression profiles (similar to doublet events in single-cell sequencing^44^).

SpatialZ enables comprehensive downstream analyses (Fig. 1e), including atlas interpolation, 3D spatial gene expression analysis, and 3D spatial domain analysis. The framework also provides *in silico* sectioning at arbitrary angles and positions, 3D rendering for direct visualization of gene expression in both volumetric and surface views, and spatial mapping of query tissue sections within the 3D atlas. Detailed methodology is provided in the Methods section.

### Evaluation on a real 3D spatial transcriptomics data

To rigorously evaluate SpatialZ’s performance in reconstructing three-dimensional tissue architecture, we utilized a publicly available STARmap dataset from the mouse visual cortex^20^. This dataset provided an ideal validation platform because it represents a real 3D spatial transcriptomics dataset generated experimentally (see Methods). We designed a systematic validation approach by artificially creating gaps in the dataset – first partitioning the 3D data into consecutive 2D sections, then holding out the even-numbered 2D sections to simulate the missing sections caused by sparse sampling in a real spatial transcriptomics experiment. This experimental design allowed us to use the held-out sections as the ground truth for evaluating the accuracy of SpatialZ’s generated virtual slices (Fig. 2a and Methods). We evaluate in terms of gene expression level, gene spatial pattern, and cell spatial localization, etc.

**Fig. 2.**
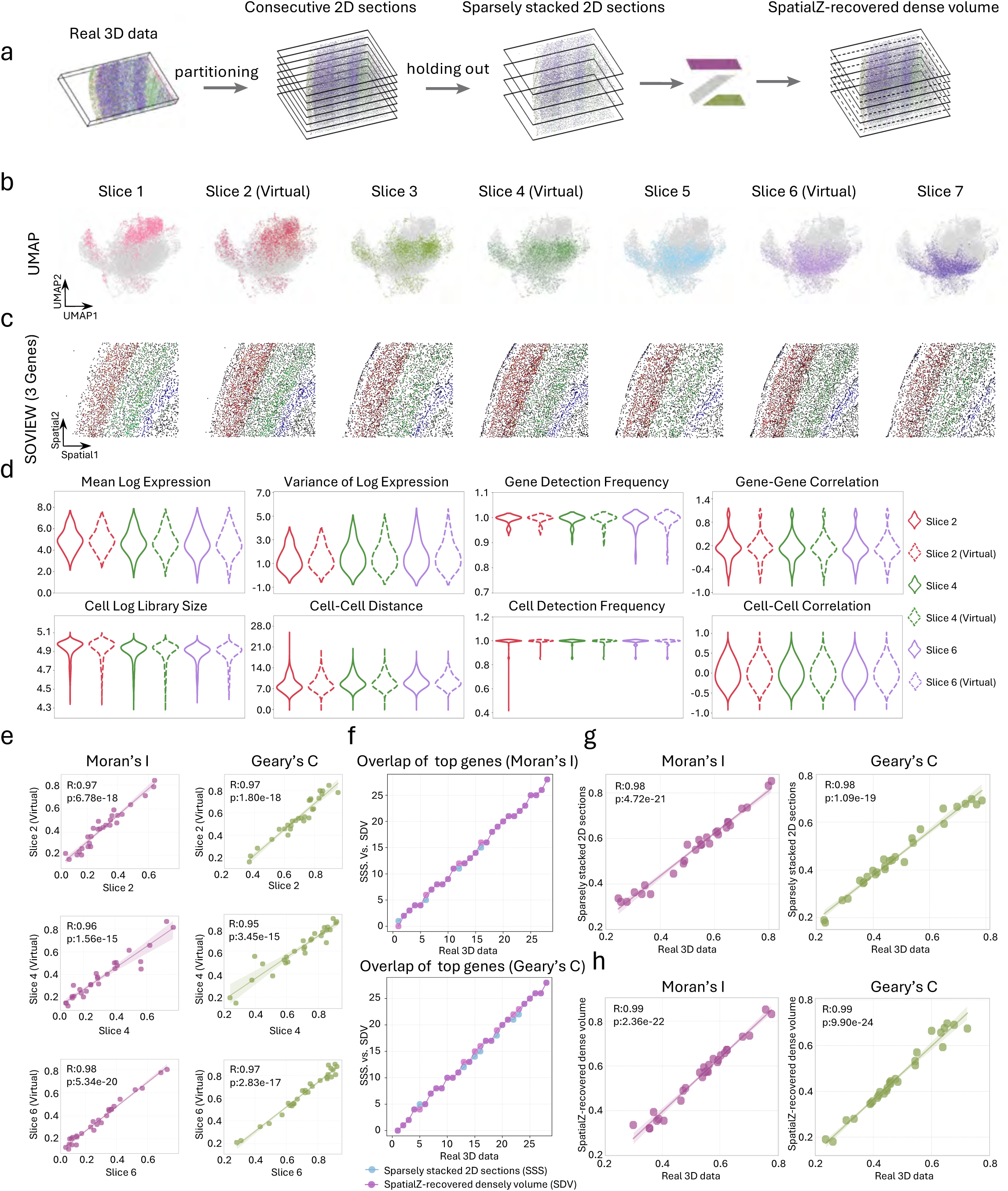
Performance evaluation of SpatialZ using real 3D spatial transcriptomics data. **a**, A real 3D data was partitioned into consecutive 2D sections and subsequently sampled as sparsely stacked 2D sections. Using the sparse data, SpatialZ recovers a dense spatial atlas by generating the leave-out intermediate sections. **b**, Uniform Manifold Approximation and Projection (UMAP) plots of real slices and virtual slices based on gene expressions in the SpatialZ-recovered dense volume. UMAP plots of the real 3D spatial transcriptomics data are presented in Extended Data Fig. 1. Volcano plots show the Kernel Density Estimation (KDE) of gene expression levels across different slices in the real 3D spatial transcriptomics data (Extended Data Fig. 2) and the SpatialZ-recovered dense volume (Extended Data Fig. 3). **c**. Spatial Omics View (SOVIEW) of three representative marker genes recovered by SpatialZ with *Cux2*, *Pcp4*, and *Flt1* indicating in red, green, and blue respectively. Spatial distributions of these marker genes are shown in Extended Data Fig. 4. Each cell was colored by gene expression level. **d**. Violin plots of eight statistic metrics of real and SpatialZ-generated virtual data.**e**. Spatial gene pattern evaluation of real and corresponding virtual slices using Moran’s I (left) and Geary’s C (right). The regression line is accompanied by a 95% confidence interval band, representing the uncertainty surrounding the regression estimate. The correlation coefficient (R) and p-value were calculated using a two-sided Spearman’s rank correlation test. **f**, Overlay of the top spatially variable genes (SVGs) between sparsely stacked 2D sections or SpatialZ-recovered dense volume versus real 3D spatial transcriptomics data in terms of Moran’s I (top) and Geary’s C (bottom). **g-h**, 3D Spatial gene pattern evaluation of comparing the sparsely stacked 2D sections (**g**) or SpatialZ-recovered dense volume (**h**) against real 3D spatial transcriptomics data. The regression line is accompanied by a 95% confidence interval band, representing the uncertainty surrounding the regression estimate. The correlation coefficient (R) and p-value were calculated using a two-sided Spearman’s rank correlation test.

To examine the gene expression landscape, we integrated all cells from both real and virtual slices, performed standard single-cell analysis and projected all cells into a low-dimensional space using Uniform Manifold Approximation and Projection (UMAP)^45,46^ (Fig. 2b). The results highlighted a continuous pattern on the gene expression manifold from slice 1 to slice 7 (Fig. 2b), consistent with the continuous pattern when using all real sections from slice 1 to slice 7 (Extended Data Fig. 1). These results indicate SpatialZ captures transitional molecular states along z-axis (Fig. 2b). The transitional gene expression states were also validated by the continuous gene expression changes along z-axis revealed by kernel density estimation (KDE) analysis on gene expression distributions from each slice along z-axis, both in all-real sections (Extended Data Fig. 2) and real-virtual slices (Extended Data Fig. 3). To examine the spatial patterns in gene expression, we visualized three layer-marker genes, *Flt1, Pcp4, and Cux2*, (Fig. 2c, Extended Data Fig. 4) using RGB color coding in the same spatial map for each section (see Methods). These genes exhibited expected laminar spatial patterns in both real and virtual slices. Other key marker genes were also validated (Extended Data Fig. 5 and Extended Data Fig. 6).

We compared the real and virtual slices quantitatively, using various evaluation metrics reflecting gene expression properties, cell properties, and spatial properties (see Methods). We found that the virtual slices generated by SpatialZ closely matched the real sections across these metrics (Fig. 2d). We additionally examined the gene expression patterns using two established statistical measures: Moran’s I and Geary’s C^47,48^. The results showed that the genes-wise spatial autocorrelation of virtual slices is significantly correlated with that of real sections (Fig. 2e). In addition, we also evaluated spatial autocorrelation in sparsely stacked 2D sections (where we artificially removed the even-numbered 2D sections in a 3D volume, representing sparse sampling in real-world scenarios) and SpatialZ-recovered dense volume (representing dense sampling generated by SpatialZ). The SpatialZ-recovered dense volume demonstrated superior performance compared to sparsely stacked 2D sections, showing more overlapping spatial genes and smaller differences in gene ranking (Fig. 2f and Extended Data Fig. 7). Correlation analysis further revealed higher consistency in the SpatialZ-recovered volume compared to the real 3D spatial transcriptomics data (Fig. 2g and 2h) (see Methods).

### Reconstruction of dense spatial cell data

To investigate SpatialZ’s capability in generating complex spatial organization at single-cell resolution, we employed an established spatially resolved cell atlas of the mouse hypothalamic preoptic region, obtained using multiplexed error-robust fluorescence in situ hybridization (MERFISH)^49^. This dataset contains a series of adjacent 2D sections. Different from previous visual cortex dataset, this dataset is more sparse (larger gap) along z-axis, so we used SpatialZ to generate 3 virtual slices between every two adjacent sections (Fig. 3a and see Methods). This dataset contains both cell type and spatial domain annotations. We examined the cell type distributions of all slices (virtual and real) on both gene expression and physical space. We also performed spatial-aware downstream tasks to test the effect of integrating the virtual slices on the performance.

**Fig. 3.**
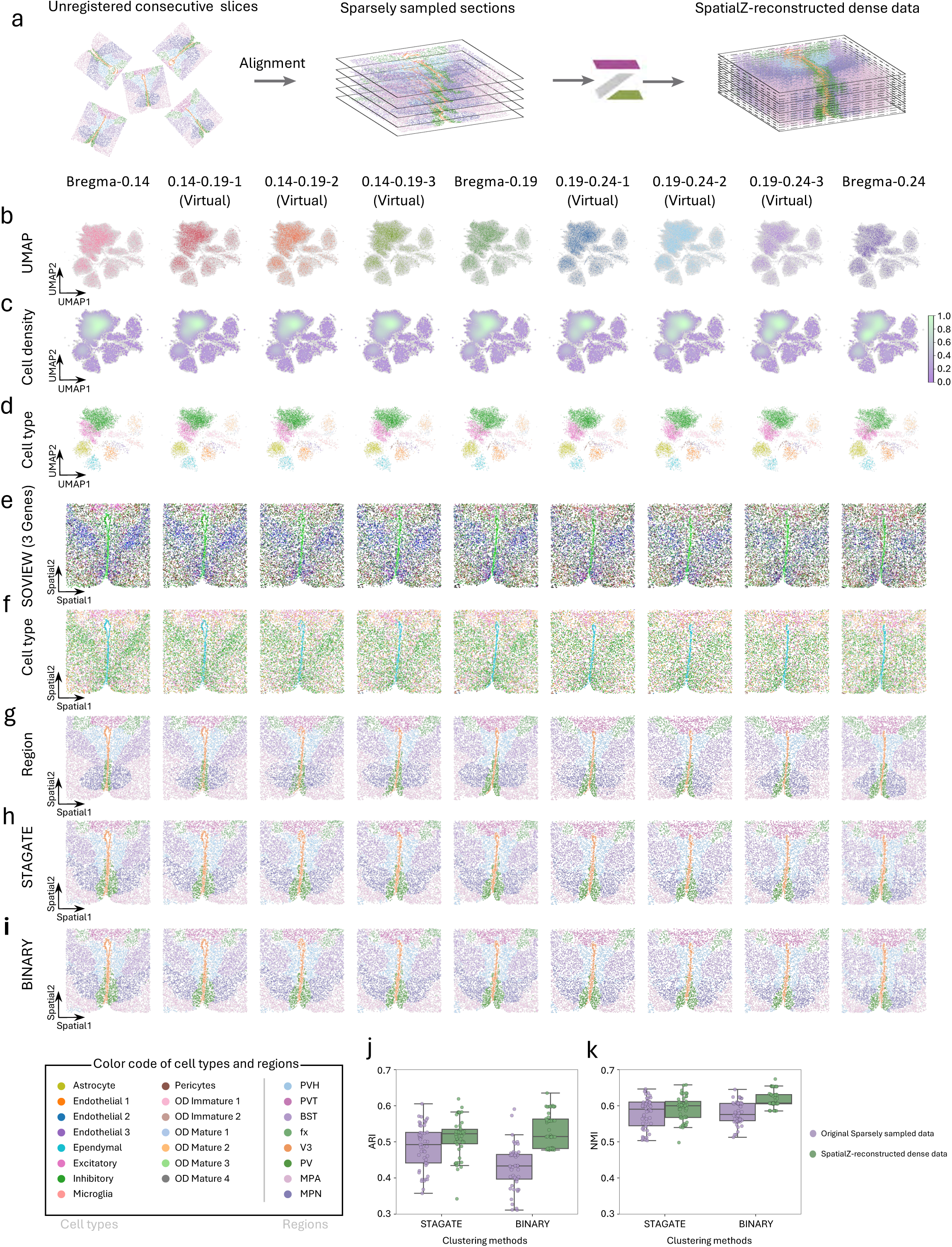
SpatialZ enables the reconstruction of a dense spatial data on complex mouse hypothalamic preoptic region. **a,** Workflow of SpatialZ for reconstructing a dense data using five single-cell spatial transcriptomics slices. **b**, UMAP plots of real and virtual slices colored by different slices in the SpatialZ-reconstructed dense spatial data. And complete UMAP plots of this SpatialZ-reconstructed dense spatial data are presented in Extended Data Fig. 8. **c**, Cell density distribution visualizations on the UMAP plots of real and virtual slices in the SpatialZ-reconstructed densely spatial data. All corresponding cell density visualizations are provided in Extended Data Fig. 9. **d**, Cell type distribution visualizations on the UMAP plots of real and virtual slices in the SpatialZ-reconstructed densely spatial data. All related UMAP plots of the cell type are presented in Extended Data Fig. 10. **e**, SOVIEW of three representative marker genes recovered by SpatialZ with *Slc17a6*, *Mlc1*, and *Gda* indicating in red, green, and blue respectively. Spatial distributions of these marker genes are presented in Extended Data Fig. 11, with each cell was colored according to gene expression level. **f**, Molecular-defined cell type visualization of both real and virtual slices under the spatial context. All spatial distribution of cell type in real slices and assigned cell type in virtual slices in the SpatialZ-reconstructed dense spatial data are showed in Extended Data Fig. 13. **g**, Transferred label visualization of both real and virtual slices under the spatial context. All spatial distribution of the ground truth in real slices and transferred labels in virtual slices are showed in Extended Data Fig. 14. Abbreviations: PVH, paraventricular hypothalamic nucleus; PVT, paraventricular nucleus of the thalamus; BST, bed nuclei of the strata terminalis; fx, columns of the fornix; V3, third ventricle; PV, periventricular hypothalamic nucleus; MPA, medial preoptic area; MPN, medial preoptic nucleus. **h-i**, Spatial domains detected by STAGATE (**h**) and BINARY (**i**). **h-i**, Spatial domain visualization of both real and virtual slices using STAGATE (**h**) and BINARY (**i**). **j-k**, Quantitative evaluation of the identified domains on the original sparsely sampling data and the SpatialZ-reconstructed densely spatial data using Adjusted Rand Index (ARI**, j**) and Normalized Mutual Information (NMI, **k**). Each point in the box plots represents an individual trial, with each trial comprising 10 runs per slice. Detailed quantitative results across the 5 slices are provided in Extended Data Fig. 16.

We integrated all cells from both real and virtual slices and performed UMAP to map every cell into the low-dimensional gene expression space. The UMAP results demonstrated remarkable consistency across all slices (Fig. 3b and Extended Data Fig. 8), with embedding density patterns showing high reproducibility between real and virtual slices (Fig. 3c and Extended Data Fig. 9). The cell type distribution on low-dimension space also showed a high consistency along z-axis (Fig. 3d and Extended Data Fig. 10).

To examine spatial gene expression patterns, we used RGB color coding scheme to visualize three region specific genes, *Slc17a6*, *Mlc1*, and *Gda*, in the same spatial map for each slice (Fig. 3e and Extended Data Fig. 11). These spatial marker genes, particularly prominent in the posterior hypothalamus, exhibited both expected spatial region patterns and gradual shifts between adjacent sections (Extended Data Fig. 12).

Complete understanding the cellular mechanisms underlying brain functions requires a detailed characterization of the spatial organization and interactions of cell types. The cellular densities and spatial distributions of cell types within virtual slices are consistent with the paired real sections (Fig. 3f and Extended Data Fig. 13). Additionally, SpatialZ enables the accurate automatic transfer of labels (such as spatial domains), eliminating the need for intensive manual annotations (Fig. 3g and Extended Data Fig. 14).

To quantitatively validate the impact of integrating virtual slices with real slices on downstream tasks, we employed two state-of-the-art spatial clustering methods - STAGATE^50^ and BINARY^51^. These methods were applied and evaluated on both the original sparsely sampled data and our SpatialZ-reconstructed dense data (Fig. 3h, 3i, Extended Data Fig. 15). The results demonstrated that the reconstructed dense spatial data achieved higher adjusted rand index (ARI) and normalized mutual information (NMI) compared to the original sparsely sampled data (Fig. 3j, 3k and Extended Data Fig. 16). These results suggested that the virtual slices can link the discontinuity of original data and benefit the downstream tasks.

### Application on a large-scale brain cell atlas

The most recent outputs of BRAIN Initiative Cell Census Network have generated hundreds of spatial gene expression sections spanning across mouse brain, allowing investigation of millions of cells within their spatial context^25,52,53^. The gap between adjacent sections motivated us to apply SpatialZ to construct a pseudo 3D atlas for mouse brain (Fig. 4a, 4b). We utilized the MERFISH brain cell atlas^52^. The dataset was registered with the Allen Mouse Brain Common Coordinate Framework version 3 (CCFv3)^52^. Given the approximately 100 μm interval between consecutive sections, we employed SpatialZ to generate nine virtual slices between each of the 129 consecutive pairs. This process resulted in a dense single-cell resolution atlas comprising 1,281 slices, containing over 38 million cells, with more than 1,100 genes profiled (Fig. 4b). This atlas currently contains the largest number of cells mapped in a mouse brain. The computational efficiency of SpatialZ enabled the construction of this comprehensive atlas in approximately 801 hours using a modest server setup (see Methods).

**Fig. 4.**
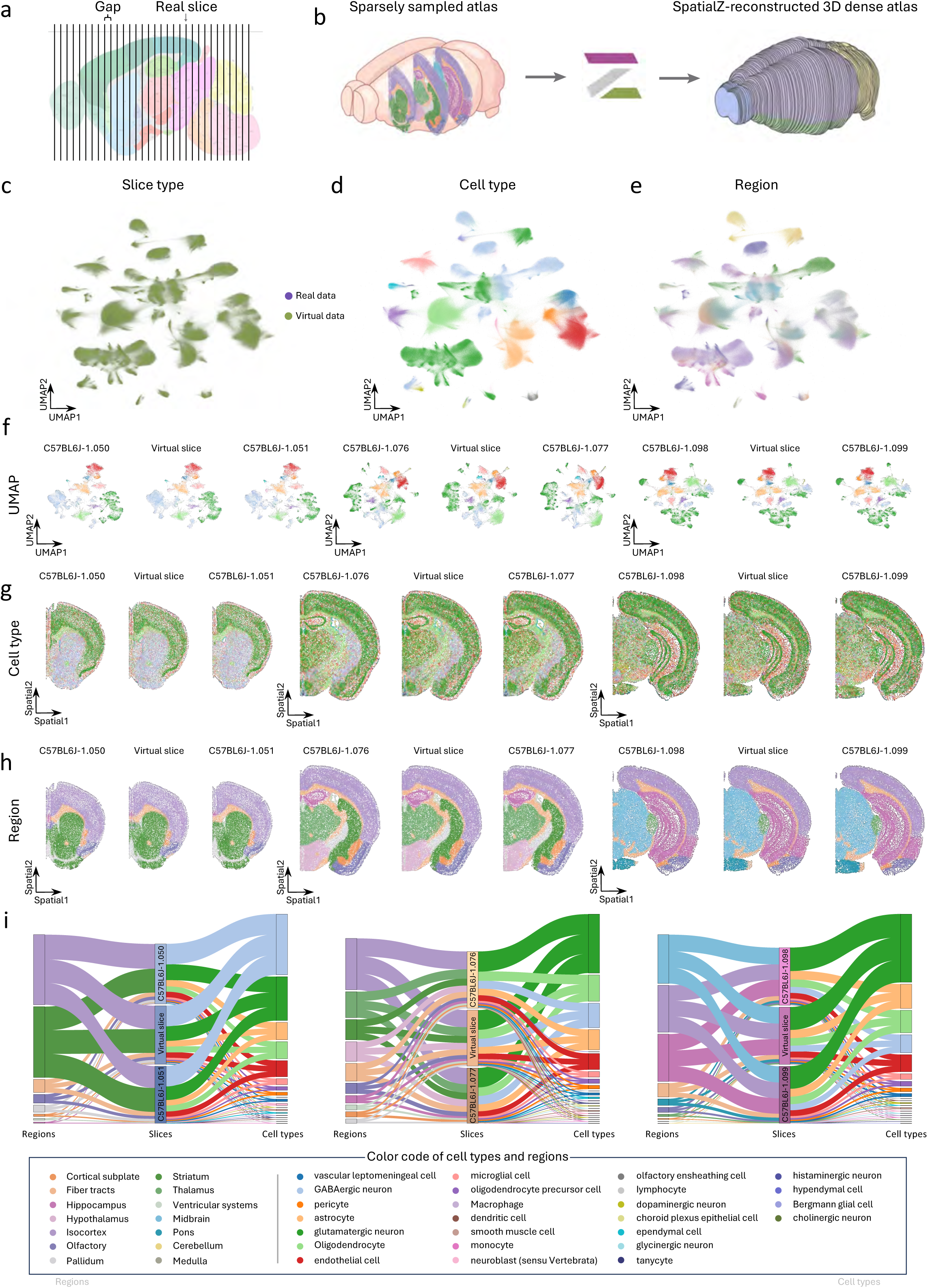
SpatialZ generates a large-scale and dense cell atlas of the mouse brain. **a,** Illustration of the brain atlas from BRAIN Initiative Cell Census Network with spatial gaps exceeding the thickness of a slice. **b**, Applying SpatialZ to bridge the spatial gaps between adjacent slices. **c**, UMAP plots of integrated real and virtual data with real and virtual data in purple and green. The number of cells from both real and virtual data was randomly downsampled at a rate of 1/10 for computational resource limitation. Detailed UMAP plots with real or virtual cells are presented in Extended Data Fig. 17a and Extended Data Fig. 17b, separately. **d**, Cell type distribution visualizations on the UMAP plots of real and virtual data. **e**, Major brain region visualization on the UMAP plots of real and virtual data. **f**, UMAP plots illustrating cell types from different regions. **g**, Cell type visualization of the real slices and virtual slice using the assigned and SpatialZ-inferred labels under spatial context. Detailed spatial distribution of cell types in real slices and virtual slices are presented in Extended Data Fig. 18. **h**, Transferred label visualization of both real and virtual slices under the spatial context. Detailed spatial distribution of the brain region in real slices and virtual slices are shown in Extended Data Fig. 19. **i**, Alluvial plots in terms of the fractions of brain regions and cell types between the intermediate virtual slice and its adjacent upper and lower slices. Detailed fractions of cell types (left) and brain regions (right) between real and virtual slices are presented in Extended Data Fig. 20.

The generated virtual slices integrate seamlessly with real sections (Fig. 4c and Extended Data Fig. 17). Both cell type labels (Fig. 4d) and transferred spatial domain labels align consistently between real and virtual slices (Fig. 4e). Given the critical importance of cell type composition and spatial organization across different brain regions, we conducted UMAP visualization of cell types from various regions of the entire mouse brain (Fig. 4f). The spatial distribution of cell types in virtual slices (Fig. 4g and Extended Data Fig. 18) and their spatial organization (Fig. 4h and Extended Data Fig. 19) showed consistent agreement with paired adjacent real sections. To validate the biological accuracy of our reconstructed atlas, we performed comprehensive analysis of cell type compositions across different brain regions. The results showed remarkable consistency between real and intermediate virtual slices (Fig. 4i and Extended Data Fig. 20).

Furthermore, we evaluated the atlas’s ability to capture known anatomical features and gene expression signatures. Key marker genes showed expected expression patterns, and cellular localization, such as cortical layering (e.g., *Cux2 and Lamp5*), hippocampus (e.g., *Prox1 and Neurod6*) and other complex structures, were accurately preserved in virtual slices (Extended Data Fig. 21). The high fidelity of these reconstructions demonstrates SpatialZ’s effectiveness in maintaining the brain spatial organization across various brain regions while maintaining biological relevance at single cell resolution.

### *In silico* sectioning and 3D gene expression rendering of mouse brain

Understanding complex three-dimensional organ structures requires examination from multiple viewing angles, yet current spatial transcriptomics technologies restrict analysis to one predefined plane - either coronal or sagittal sections typically^52,54^. This limitation exists because a biological brain cannot be sectioned and assayed in coronal and sagittal planes simultaneously. Such constraints significantly impact our ability to comprehend intricate spatial relationships and molecular gradients that might be more apparent from alternative viewing angles. While it is theoretically possible to generate 2D spatial atlases in multiple planes using different animals, this approach faces substantial challenges. The considerable costs, technical difficulties, and reproducibility concerns (since different animals are used) make such comprehensive sampling impractical. SpatialZ offers an alternative solution that enables multiple analytical perspectives from a single specimen.

We developed an “*in silico* sectioning” module to generate synthesized 2D slices from any angle using the SpatialZ-reconstructed 3D atlas (Fig. 5a left, see Methods). Using the pseudo 3D atlas derived from the coronal atlas, we performed *in silico* sectioning to synthesize 50 sagittal slices and 50 horizontal slices spanning the entire brain hemisphere. Furthermore, we generated 50 oblique slices (custom angle) at 45-degree angles relative to conventional planes, providing unprecedented flexibility in analyzing brain organization (Fig. 5b and Methods). To validate the biological consistency of our synthesized data, we performed comprehensive analysis of cell type compositions across different brain regions, revealing continuous molecular gradients and spatial relationships (Extended Data Fig. 22). This multi-perspective capability allows researchers to examine tissue architecture from any desired angle without requiring additional experimental data collection. To validate the biological accuracy of synthesized slices, we additionally compared them systematically with the Allen Brain Atlas in situ hybridization (ISH) reference data. We examined five key marker genes (*Slc17a7*, *Slc30a3*, *Penk*, *Gfap*, and *Agt*) across different brain regions in both sagittal and coronal planes (Fig. 5c and Extended Data Fig. 23).

**Fig. 5.**
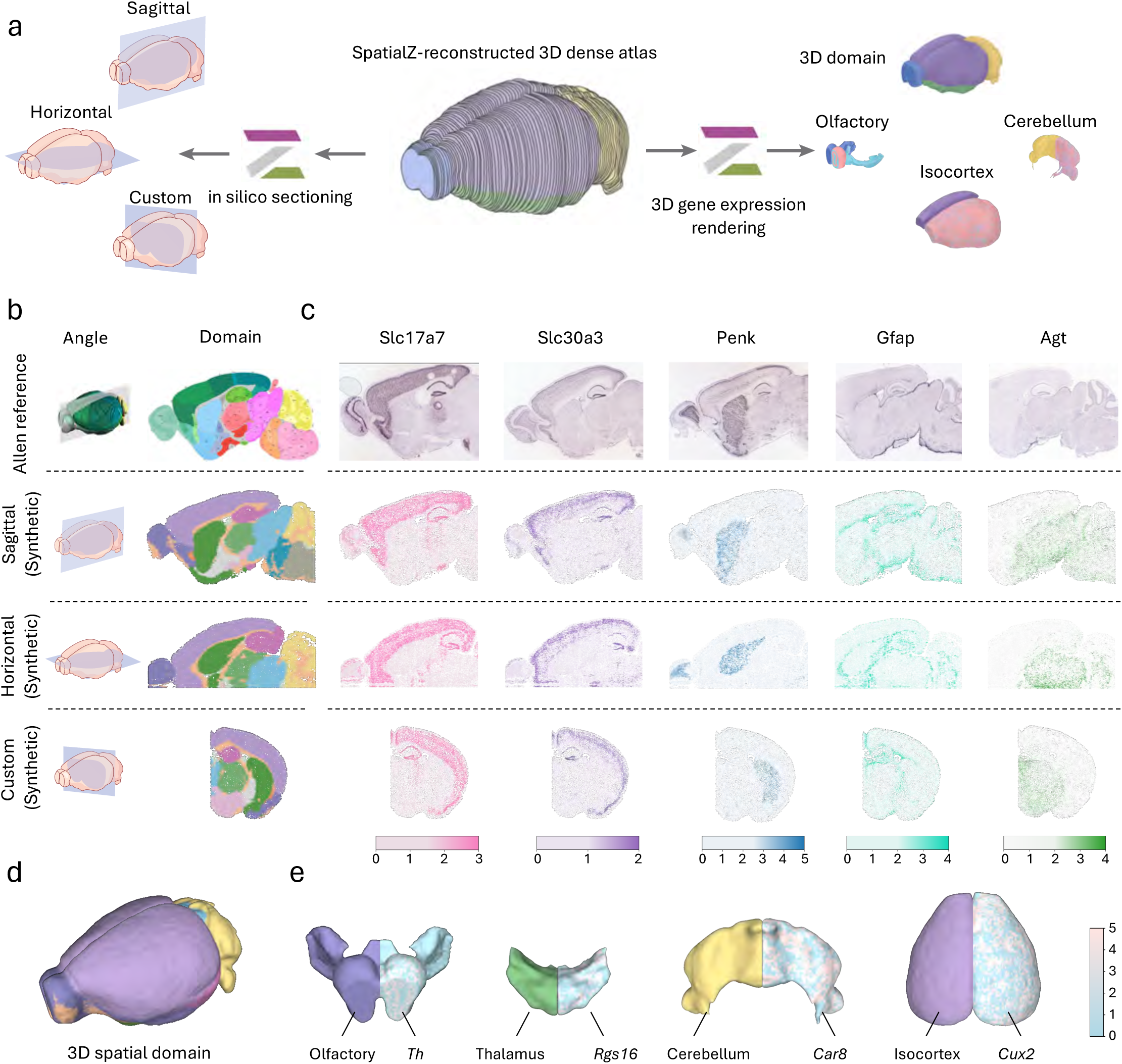
SpatialZ supports *in silico* sectioning and 3D gene expression rendering. **a,** Schematic of *in silico* sectioning (left) and 3D gene expression rendering (right) using a SpatialZ-reconstructed 3D spatial atlas regarding the coronal planes.**b**, Visualization of plane angles and brain regions on Allen reference and our SpatialZ- synthesized new different view planes (sagittal, horizontal, and custom). The first row shows the Allen reference with a sagittal angle and corresponding anatomical reference atlas. The second, third and fourth rows depict sagittal, horizontal, and custom planes, respectively. The custom plane in our experiment is the oblique angle with 45-degree angles relative to conventional planes. **c**, Marker gene visualization across different planes (sagittal, horizontal, and custom) under the spatial context, with cells colored by gene expression levels. The first row shows reference gene expression patterns from the Allen Brain Atlas in the sagittal plane. **d**, 3D continuous mesh rendering of mouse brain colored by major brain regions. **e**, Visualization of the 3D gene expression gradients of different regions. The left side and right slide of each illustration show the 3D structure of a specific mouse brain region (from left to right: Olfactory, Thalamus, Cerebellum, Isocortex) and the corresponding marker gene, respectively (from left to right: *Th*, *Rgs16*, *Car8*, *Cux2*).

To enhance three-dimensional data visualization and analysis further, we implemented a “3D Continuous Tissue Rendering” module for SpatialZ (Fig. 5a right, 5d). This module transforms discrete cellular data into continuous three-dimensional meshes using advanced interpolation algorithms (see Methods). The rendering system maintains spatial accuracy while providing smooth transitions between adjacent regions, enabling clearer visualization of complex spatial relationships that may be unclear in traditional visualization approaches (Fig. 5e and Extended Data Fig. 20). The continuous rendering capability enables advanced analytical features, including visualization of gene expression gradients, interactive exploration of cellular organization, quantitative analysis of spatial relationships, integration of multiple molecular markers, and identification of region-specific 3D gene expression features.

The combination of synthesized views and continuous rendering capabilities represents a significant advancement in spatial transcriptomics analysis. These tools enable researchers to examine tissue architecture from any view, identify previously obscured spatial patterns, and analyze molecular gradients across three-dimensional space. To facilitate adoption, we have made the complete toolkit freely available through our open-source software package. This comprehensive approach, coupled with SpatialZ’s dense reconstruction capabilities, provides researchers with powerful new tools for investigating spatial gene expression patterns and cellular organization in complex tissues, while maintaining biological accuracy and analytical flexibility.

### The SpatialZ-reconstructed 3D atlas provides a robust reference for precise spatial mapping of query samples

Recently, an advanced method called cross-sample alignment of spatial omics (CAST) has been developed by Xiao’s group to enables spatial mapping of tissue sections^21^. But its original demonstration is limited to 2D analysis. We enhanced this capability by integrating CAST with SpatialZ, creating a powerful system for three-dimensional sample searching and locating (Fig. 6a).

**Fig. 6.**
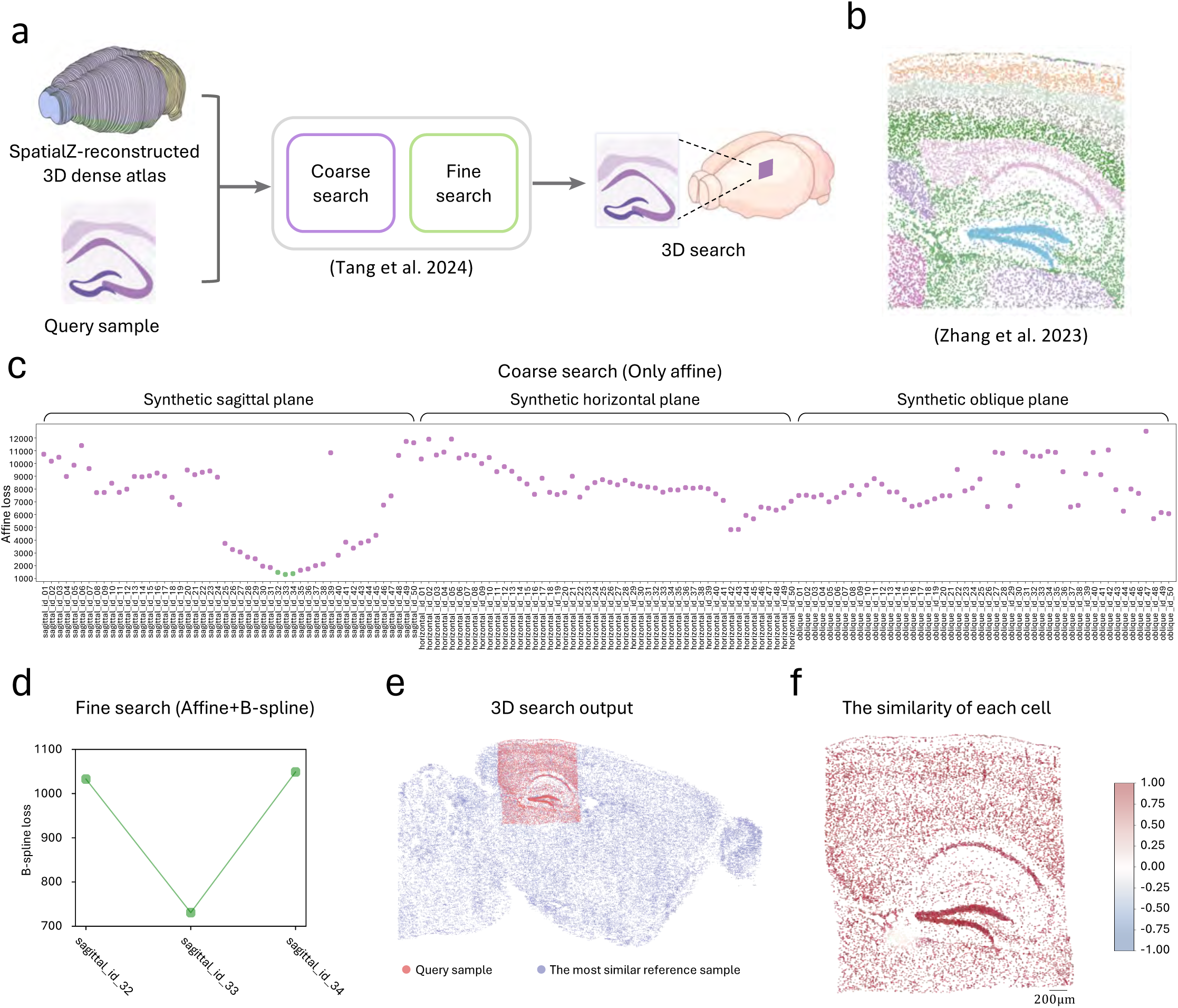
The SpatialZ-reconstructed 3D atlas provides a robust reference for accurate 3D spatial search. **a**, Schematic demonstration of 3D spatial search using the SpatialZ- reconstructed 3D atlas. We integrated CAST with SpatialZ for three-dimensional sample searching and locating. The 3D spatial search includes two processes: coarse search (quick rigid alignment) and fine search (finer nonrigid alignment). **b**, The query sample consists of a truncated tissue section encompassing the hippocampus and a partial cortical region, derived from a sagittal slice and measured using MERFISH (Multiplexed Error-Robust Fluorescence In Situ Hybridization). **c**, Scatter plot illustrating affine loss between query sample and reference sample within SpatialZ-reconstructed 3D atlas during coarse search phase. To optimize computational efficiency, 150 synthesized slices (50 sagittal, 50 horizontal, and 50 oblique) were utilized as a 3D multi-perspective reference atlas. The lowest three green dots (in sagittal planes) indicate the most probable matches for query sample, providing potential candidates for subsequent fine alignment. **d**, Final match result after fine search process. The query sample was localized to the location of the 33rd slice of the sagittal plane where represents best alignment of the specific anatomical location. **e**, Visualization of 3D spatial search output between the query sample and the 33rd slice of the sagittal plane in SpatialZ- reconstructed 3D atlas. **f**, The similarity of each cell in the query sample with the 33rd slice of the sagittal plane. Each cell in the query sample is colored by adjusted Pearson correlation.

To demonstrate this capability, we used a truncated hippocampus tissue sample, comprising 13,853 cells from an experimentally obtained mouse sagittal brain section, as our query sample (Fig. 6b). For the reference, we created the data by generating sectional views from three distinct angles using our previously described *in silico* sectioning method. More planes could be used if needed. The mapping process involves two steps: First, we performed a rough alignment between the query sample (S1) and the multi-angle references through affine transformation (Fig. 6c). This preliminary analysis identified the most similar spatial orientations and section positions within the 3D reference atlas that match the query sample (green dots in the scatter plot indicate the highest similarity) (Fig. 6c). Specifically, the three sections most closely matching the query sample were the 32nd, 33rd, and 34th synthetic sagittal sections. Second, we conducted a more refined search using B-spline free-form deformation (FFD), which identified the 33rd synthetic sagittal section as the optimal match (Fig. 6d). The high correlation between each query cell and its nearest reference cell (Fig. 6e, 6f) confirms the accurate alignment and precise localization of the query sample within the SpatialZ-generated 3D reference atlas.

This successful integration of CAST and SpatialZ demonstrates the feasibility of extending traditional 2D mapping to comprehensive 3D analysis. While we used three angles for this demonstration, the method can accommodate additional angles and sections for more precise localization in practical applications.

## Discussions

SpatialZ represents a significant advancement in spatial transcriptomics analysis, addressing fundamental limitations in current 3D tissue analysis approaches. While existing technologies are restricted by high cost, complex protocols, and substantial gaps between tissue sections, our framework uniquely generates intermediate slices between sparsely sampled sections, creating comprehensive pseudo-3D atlases. This capability particularly addresses critical bottlenecks in developing spatial foundation models and whole-organ mapping, where experimental limitations and data availability often constrain progress.

The framework’s ability to generate numerous virtual slices advances spatial transcriptomics by expanding datasets with biologically plausible sections, enabling comprehensive organ-level analysis previously impossible through experimental methods alone. Our validation using real 3D spatial transcriptomics data and independent 2D serial sectioning demonstrated strong consistency between virtual and real slices, confirming the framework’s reliability for complex tissue analysis.

However, several key limitations warrant acknowledgment. First, SpatialZ generates a pseudo-3D atlas through interpolation, with quality dependent on initial 2D slice data spacing and quality. Integration with emerging real 3D imaging technologies and implementation of tissue-specific adaptive interpolation algorithms could address these limitations. Second, the framework’s performance depends on input quality and can be affected by tissue processing artifacts. Future versions will incorporate machine learning approaches for automated artifact detection and correction, alongside robust quality control pipelines^55^.

Despite these limitations, SpatialZ’s impact extends beyond spatial transcriptomics by enabling comprehensive multi-species and multi-organ spatial atlases. The framework’s demonstrated ability to process over 38 million cells efficiently, generate arbitrary section views, and enable continuous visualization of spatial molecular expression profiles provides researchers with powerful tools for investigating tissue organization and function, particularly valuable where traditional approaches are limited by technical or practical constraints.

In conclusion, while acknowledging current limitations, SpatialZ represents a significant step toward understanding complex biological systems through spatial transcriptomics, bridging the gap between 2D and 3D analysis. Future development will focus on addressing limitations through technological advancement, improved validation methods, and adjustment for temporal data interpolation. This comprehensive approach opens new avenues for investigation in life sciences in 3D.

## Methods

### Data processing

In our experimental setup, we utilized several sets of publicly available spatial transcriptomic data to evaluate SpatialZ’s performance in reconstructing three-dimensional tissue architecture. We first utilized a publicly available STARmap dataset (Data 1) from the mouse visual cortex^20^. This dataset served as an ideal validation platform, representing a real 3D spatial transcriptomics dataset generated experimentally. We filtered out the uppermost (z-coordinates 6 to 13) and lowermost layers (z-coordinates 91 to 94) from the real 3D data to avoid technical noise and unreliable areas. We then segmented the remaining tissue into seven consecutive sections, removing sections 2, 4, and 6 to create sparsely stacked 2D sections. The remaining sections (1, 3, 5 and 7) were used to sequentially infer the missing sections (virtual slices 2, 4, and 6) for subsequent performance comparison and spatial analysis. To explore SpatialZ’s capability in generating complex spatial organization at single-cell resolution, we used a spatially resolved cell data from the mouse hypothalamic preoptic region obtained using MERFISH technology (Data 2)^49^. This dataset, characterized by larger spatial gaps, includes both cell type and spatial domain annotations. We employed SpatialZ to generate three virtual slices between every two adjacent sections and examined gene expression, cell type distributions and label transfer across all slices (virtual and real). Lastly, to validate SpatialZ’s applicability to a large-scale brain cell atlas, we leveraged data from the BRAIN Initiative Cell Census Network (Data 3)^52^. This data set provided hundreds of spatial gene expression sections across the mouse brain. Using SpatialZ, we interpolated and generated nine virtual slices between each of the 129 consecutive real pairs, constructing a complete 3D hemisphere atlas.

### Spatial alignment

To generate accurate spatial locations for virtual slices, it is essential to align the sparsely sampled sections into a common coordinate system. For the data (Data 3) with a pre-registered coordinate system, we align sections using the coordinates from the registered system, such as the Allen Mouse Brain Common Coordinate Framework version 3 (CCFv3)^43^. For data (Data 2) without any pre-registered coordinate system, we use PASTE^56^, which integrates gene expression similarity and physical location proximity between cells in adjacent slices.

### SpatialZ algorithm

Given a sequence of sparsely sampled sections from the real tissue, the input for each slice *slice*_*k*_ includes gene expression matrix ***X***^*k*^ ∈ ℝ^*N*_*k*_×*M*^, where *N*_*k*_ indicates the number of cells in k-th section and *M* denotes the feature dimension of cells (e.g., gene expression panel size), corresponding registered 2D spatial coordinates ***S***^***k***^ ∈ ℝ^*N*_*k*_×2^ and the molecularly defined cell types **C**^*k*^ ∈ ℝ^*N*_*k*_^, SpatialZ will sequentially generate virtual slices based on these sparsely sampled sections.

#### Location generation

To accurately generate the spatial positions of cells within the interpolated virtual slice, we model the 2D spatial distribution of cells in adjacent slices and their spatial gradient along the z-axis. We expedited the learning process by employing a combination of sampling strategies to initialize a composite set of cellular locations. The value of in the adjacent slices are denoted by ***S**_k_* ∈ ℝ^*N*×2^ and ***S**_k_*_+1_∈ ℝ^*N*_*k*+1_×2^, the generated cellular locations within the virtual slices is represented as ***S**_vir_* ∈ ℝ^*N_vir_*×2^, where *N_vir_* is the number of generated cells within the virtual section. The value of *N_vir_* can be determined in two ways: (1) a manually defined value, *N*_*custom*_, or (2) an automatically estimated value, *N*_*estimate*_, which is derived from the distribution of cell counts in the adjacent slices: 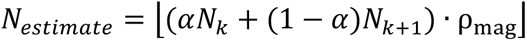

where *α* ∈ [0,1] is the proximity coefficient that controls the relative weights between upper and lower slices, and ρ_mag_ is a magnification factor to adjust the density of the synthesized slice, default is 1. The symbol ⌊·⌋ denotes the floor operation. Next, we uniformly sample ⌊*αN*_*k*_⌋ coordinates from **S**_*k*_ and ⌊(1 − *α*)*N*_*k*+1_⌋ coordinates from ***S***_*k*+1_ ∈ ℝ^*N*_*k*+1_×2^. The two sampled sets are then concatenated to form the initial coordinate set 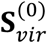.

To further optimize the generated cellular location within the interpolated virtual slices, we minimized the Sliced Wasserstein Distance (SWD) to refine the spatial distribution. The loss function is defined as:

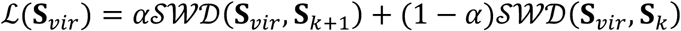

We utilize a Monte-Carlo approximation method to compute the *SWD_p_*(*μ*, *v*):

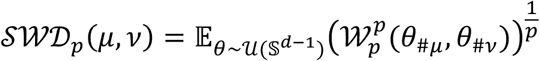

where *θ*_#*μ*_ and *θ*_#*v*_ represent the pushforwards of the projection **S**_*vir*_ ∈ ℝ^*d*^ ↦ 〈*θ*, ***S***_*vir*_〉, and 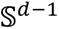 denoted the unit sphere in ℝ^*d*^. The parameter *p* refers to the p-norm used when computing the Wasserstein distance, with a default value of 2. We used stochastic gradient descent to minimize this loss function, updating the coordinate set **S**_*vir*_ as follows:

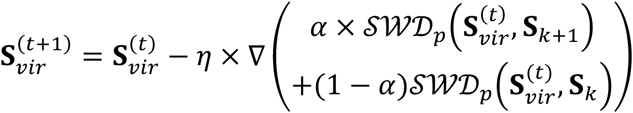

where *η* is the learning rate, ∇ represents the gradient of 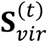, and *t* represents the iteration time of the optimization. Through iterative updates, we optimize **S**_*vir*_ to ensure that interpolated virtual slice align with the spatial distribution of the adjacent sections while maintaining smooth continuity in the 3D space.

#### Attribute assignment

We assign cell types and other cell attributes (e.g., region labels) for synthesized cells by using the k-nearest neighbors (k-NN) algorithm, where the Euclidean distance is used to compute the nearest neighbors from two real adjacent sections:

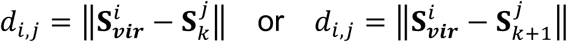

For each synthesized cell, we first identify *k*_*ct*_ nearest neighbors in two adjacent slices, forming nearest neighbor set Ι_1_[*i*] and Ι_2_[*i*], respectively. The cell type distribution within the virtual slice is the spatial distance weighted distribution from Ι_1_[*i*] and Ι_2_[*i*], the weight can be calculated by:

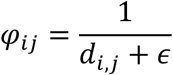

where ϵ is a small constant to prevent division by zero when the distance *d*_*i,j*_ is very small. Finally, the cell type *c*_*i*_ for synthesize cell is determined using the following formula:

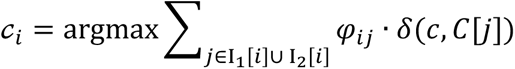

Here, *δ*(*c*, *C*[*j*]) is an indicator function that equals 1 if the cell type *c* matches the cell type *C*[*j*] of the neighbor, and 0 otherwise.

After assigning cell types to all synthesized cells, we performed a reverse lookup to identify the nearest cell with the same cell type. Label information (e.g., spatial domain labels, if available) from the neighboring cell was then transferred to the corresponding synthesized cell. This step ensures the accuracy of the spatial and functional context of the synthesized cells, based on the characteristics of their closest real counterparts.

#### Gene expression synthesis

Considering that cellular gene expression is influenced not only by the intrinsic cell type but also by the surrounding spatial context. To capture this complexity, we employed Multi-range Cell Context Decipherer (MENDER)^57^ to capture the spatial niche for each cell. MENDER provides a structured and vectorized way to represent the local cellular context across multiple scales. Other related embedding methods can also be employed^58–60^. Specifically, let *N*_*k*_ be the total number of cells in *k*-th slice, *C* be the number of unique cell states (or types), and *S* be the number of predefined spatial niche scales. For a given cell *i* in *k*-th slice, the spatial niche **E**_*i*,*k*_ ∈ ℤ^*S*×*C*^ records the count of neighboring cells within each range *s* that belong to each cell state *τ*_*c*_:

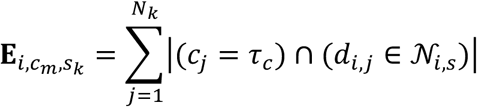

where *j* represents the neighboring cells of cell *i*, *d*_*i,j*_ represents the Euclidean distance between cells *i* and *j*, and *τ*_*c*_ represents a specific cell type. *N*_*i*,*s*_ = [(*s* − 1) × ℛ, *s* × ℛ), ℛ is a fixed radius increment. Consequently, the spatial niche for cell *i* is represented as:

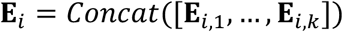

To quantify the similarity of spatial niche representation between cells, we use cosine similarity to assess the similarity of the spatial niche between cells in the same states. Given two cells *i* and *j*, with spatial niche representations **E**_*i*_ and **E**_j_, the cosine similarity *S*(*i*, *j*) was calculated as follows

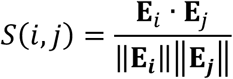

where ||**E**_**i**_|| and ||**E**_**j**_|| denote the Euclidean norms of **E**_*i*_ and **E**_j_, respectively. The cosine similarity *S*(*i*, *j*) measures the degree of similarity between cells *i* and *j* based on their microenvironmental context, with higher values indicating greater similarity.

Lastly, gene expression profiles for synthesized cell are generated based on spatial niche similarity through probabilistic inference. Specifically, for each synthesized cell *i* with assigned cell type *c*_*i*_, we first identified its *k*_sam_ spatial nearest neighbors 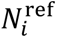 from its adjacent sections, considering cells of the same type:

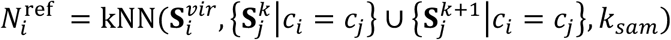

Next, the precomputed spatial niche similarities are weighted by the spatial nearest neighbors 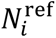:

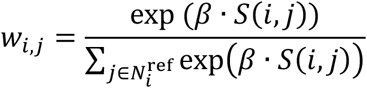

where *β* is a scaling factor, with a default value of 100. The expression value for gene *g* in synthesized cell *i* **X**_*i*_[*g*] is obtained by probabilistically sampling from the reference cells, with the probability proportional to *w*_*i,j*_. Mathematically,

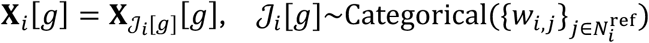

where 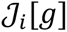 is a random index that represents the reference cell selected for gene *g*, drawn from a categorical distribution with probabilities 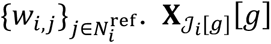 is the expression value of gene *g* from the selected reference cell.

### *In silico* sectioning

The “*In silico* sectioning” module is designed to synthesize a novel view from a SpatialZ- reconstructed 3D atlas along any specified angle (such as sagittal, horizontal, even custom plane). Users simply need to input the desired rotation angle and displacement offset into the “*In silico* sectioning” module and SpatialZ will automatically rotate the coordinate system, apply the specified offset, and synthesize the new sectional data from the chosen view.

#### Coordinate system rotation

To synthesize a sectional view at any desired angle, the first step is to rotate the coordinate system of the SpatialZ-generated 3D atlas. Let **S** ∈ ℝ^*N*×3^ be the cell coordinates for the 3D atlas, where *N* is the number of all cells, and **S**_*i*_ = [*x*_*i*_, *y*_*i*_, *z*_*i*_] represents the 3D coordinates of each cell. We calculate the geometric center **c**_**center**_ ∈ ℝ^3^ as follows:

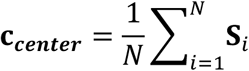

The centered coordinates matrix **S**_*centered*_ ∈ ℝ^*N*×3^ is then obtained by subtracting the geometric center from each cell:

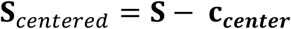

Next, to rotate the 3D atlas to the desired slicing angle, we define rotation matrices for each of the x, y, and z axes. The rotation matrices are separately given by:

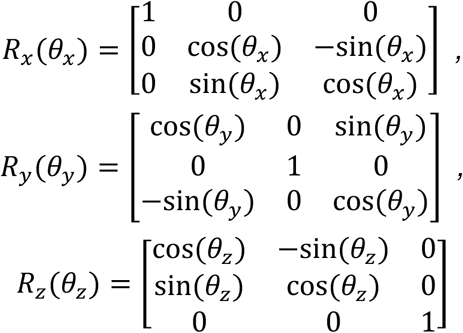

where *θ*_*X*_, *θ*_*Y*_ and *θ*_*Z*_ represent the angle of rotation about the x, y and z axises, separately.

To achieve the desired arbitrary angle of slicing, we combine three rotation matrices to form the 3D composite rotation matrix *R*_3*D*_:

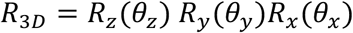

The centered coordinate matrix **S**_*centered*_ is then rotated using *R*_3*D*_, resulting in the rotated coordinates **S**_rcdadee_ ∈ ℝ^3^:

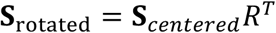

Each cell the rotated coordinate system is denoted by **S**_rcdadee,*i*_ = [*X*_rcdadee,*i*_, *Y*_rcdadee,*i*_, *Z*_rcdadee,*i*_].

#### Section position determination

Once the coordinate system has been rotated, the next step is to determine the position of the synthesize slice. We determine the center along the *Z* axis in the rotated coordinate system. The range of *Z* values after rotation is given by [*Z*_*min*_, *Z*_*max*_]:

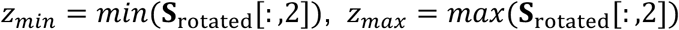

The position of the slice *Z*_*slice*_ is determined by taking the midpoint of the *Z* axis range and adding an offset *a*_*offset*_ to adjust the slice position along the *Z* axis:

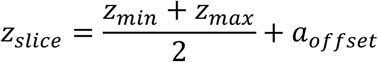

Here, *a*_*offset*_ is a parameter used to adjust the slice position along the *Z* axis.

#### New view synthesis

To synthesize new view section data, we extract the relevant cells on the predetermine slice and projected them onto a two-dimensional plane. The thickness of the slice defines the range of *Z* values to be included in the slice. Given the slcie thickness *d*, the cells that belong to the slice are those that satisfy the condition:

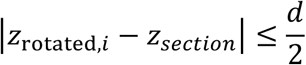

Finally, the two-dimensional projection coordinates are defined as follows:

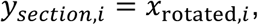

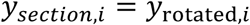

### 3D Continuous Tissue Rendering

#### 3D spatial domain point cloud generation

We start by utilizing the SpatiaZ-reconstructed atlas to extract spatial cell coordinates (x, y, z) along with corresponding spatial labels (e.g., brain region labels) to construct a global point cloud representation of distinct spatial domain. The resulting point clouds, which include both coordinate information and the color properties of each cell, are then stored in PLY file format for further analysis and visualization.

#### Field of view-specific point cloud generation

Different with the global point cloud generation, this process focuses on creating localized representations by selecting a specific field of view (FOV) of interest. This process begins by filtering the spatial coordinates and associated labels to target only a particular region, effectively narrowing down the data to the area of interest. This enables a detailed examination of finer spatial features and structures within a targeted domain, which may not be evident in the global representation. The resulting point cloud is stored in PLY format for detailed visualization and deeper region-specific analysis.

#### spatial gene expression point cloud generation

We extended our approach to capture spatial gene expression patterns by specific genes of interest, extracting their corresponding expression levels alongside spatial coordinates. By integrating both spatial information and gene expression data, we create a point cloud representation that highlights how particular genes are expressed across different regions. By extracting the cells that express the target genes, we use their spatial coordinates to construct a 3D point cloud. Gene expression intensity is visualized through color mapping. This enriched representation serves as tool for analysising localized gene functions and variations. As in previous steps, the resulting point cloud is stored in PLY format for the analysis and visualization of the spatial distribution of gene expression.

#### 3D continuous mesh construction

Building upon the previously generated point cloud, this step utilizes Open3D’s meshing algorithms to transform scattered point cloud data into a smooth, continuous mesh. First, the point cloud is downsampled using voxel_down_sample() to accelerate mesh generation process while preserving the overall geometry. Next, denoising is performed using remove_statistical_outlier(), which removes noisy points by analyzing the neighborhoods of each point. The denoising process is repeated twice to ensure a clean point cloud for reconstruction. Once the point cloud is cleaned, surface reconstruction is performed using the Alpha Shape algorithm via create_from_point_cloud_alpha_shape(). The alpha parameter is set to 300 to control the level of detail, ensuring the generation of a coherent mesh. Vertices and colors from the reconstructed mesh are then extracted to create a new point cloud using surface_point_cloud(). Normals are estimated using the kNN algorithm via estimate_normals(), with knn set to 100 to ensure smooth and reliable normal estimation and using orient_normals_consistent_tangent_plane() to ensure the same direction normal. To generate a continuous surface, Poisson reconstruction is applied using create_from_point_cloud_poisson(), with octree depth parameter is set to 9, balancing mesh quality and computational efficiency. The mesh is then smoothed with filter_smooth_simple() over five iterations to remove small-scale noise while preserving the overall structure. To enhance visualization, KDTreeFlann() is used to maintain visual consistency with original data. Finally, the generated mesh is then visualized and further refined using MeshLab^61^, a software as known for its robust mesh processing capabilities. The result output is a highly detailed and continuous mesh, offering an insightful and comprehensive visualization of tissue architectures.

### 3D spatial search algorithm

We utilize CAST^21^, a cross-sample alignment method for spatial omics data, to search and match a real and truncated sagittal sample, encompassing the hippocampus and part of cortical region, against our large-scale SpatialZ-reconstructed atlas. We integrated CAST with SpatialZ to create a powerful system for three-dimensional sample searching and locating. During the first phase, an affine transformation was used to coarsely search and match the three most similar slices, characterized by the lowest CAST stack loss values, from 150 synthesized slices (comprising 50 sagittal, 50 horizontal, and 50 oblique slices). In the second phase, we accurately locate the query sample by B-spline free-form deformation (FFD). The SpatialZ-reconstructed atlas proved to be a comprehensive reference resource, supporting precise spatial alignment and 3D spatial search.

### Evaluation metrics

#### Gene and cell level summary statistics

In Figure 2d, we present a comparative analysis of the distributions of eight distinct summary statistics calculated at four different levels: gene, cell, gene-pair, and cell-pair levels, comparing both real and virtual slices. The summary statistics are defined as follows: (1) Mean of Log Expression (Gene-Level): This metric represents the mean of the log-transformed expression values for a given gene, aggregated across all cells. (2) Variance of Log Expression (Gene-Level): This statistic quantifies the variance of the log-transformed expression values for each gene, calculated across all cells. (3) Gene Detection Frequency (Gene-Level): This metric describes the detection rate of a given gene, measured as the proportion of cells in which the gene has nonzero counts. (4) Gene-Gene Correlation (Gene-Pair-Level): This statistic captures the correlation between the log-transformed expression values of two genes across all cells. (5) Cell Log Library Size (Cell-Level): This represents log-transformed total read or unique molecular identifier (UMI) count for a cell. (6) Cell-Cell Distance (Cell-Pair-Level): This measure refers to the Euclidean distance between two cells, calculated in a 50-dimensional principal component space derived from the cell-by-gene log-transformed matrix. (7) Cell Detection Frequency (Cell-Level): This value reflects the detection frequency of a given cell, computed as the proportion of nonzero counts across all genes. (8) Cell-Cell Correlation (Cell-Pair-Level): This correlation is computed between two cells’ log-transformed expression values, aggregated across all genes.

#### Evaluation metrics for spatial gene pattern

To evaluate the spatial expression patterns between real and virtual slices, we utilized Moran’s I and Geary’s C as quantitative measures of gene expression spatial autocorrelation. Moran’s I is used to assess the overall spatial autocorrelation within a dataset, and its formula is given by:

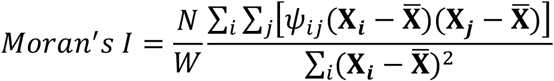

where **X**_**i**_ and **X**_**j**_ are the gene expression values at cell *i* and *j*, respectively, 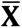 represents the mean gene expression, *N* is the total number of cells, and ψ_*i*j_ represents the spatial weight between cells cell *i* and *j*. Another metric commonly used for assessing spatial autocorrelation, provides a measure focused on local differences. Geary’s C is defined as:

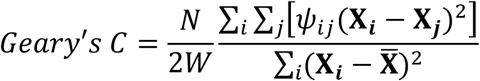

#### Evaluation metrics for spatial domain

To evaluate the spatial domain performance compared to the original sparsely sampled atlas and the SpatialZ-reconstructed dense spatial atlas, we utilize two key metrics: Adjusted Rand Index (ARI) and Normalized Mutual Information (NMI). The Adjusted Rand Index (ARI) measures the degree of agreement between manual labels and spatial clustering results. ARI values range from −1 to 1, with 1 indicating perfect agreement, and values near 0 suggesting clustering results are no better than random assignment. The ARI is computed as follows:

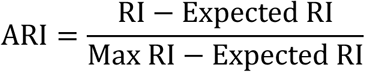

where the RI is the Rand Index, defined as:

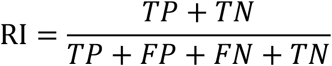

where *TP* (True Positive) refers to pairs of samples that are correctly grouped into the same cluster both in the clustering results and the true labels, *TN* (True Negative) refers to pairs correctly assigned to different clusters, *FP* (False Positive) denotes pairs incorrectly placed in the same cluster, and *FN* (False Negative) denotes pairs incorrectly placed in different clusters. Another metric used is Normalized Mutual Information (NMI), which measures the shared information between clustering results and reference labels, normalized by the average entropy of each distribution. NMI values range from 0 to 1, with 1 indicating perfect correlation, meaning clustering perfectly predicts the true labels. NMI is calculated as:

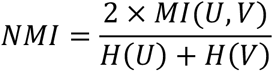

where *MI*(*U*, *V*) is the mutual information between the cluster assignments *V* and the true labels *U*, *H*(*U*) is the entropy of the true labels, and *H*(*V*) is the entropy of the cluster labels.

### SOVIEW

SOVIEW is a powerful visualization and analysis tool of spatial omics data, enabling effective inspection of genes’ spatial expression^62,63^. We utilized SOVIEW to visualize the inherent continuity within tissue structures and to examine the distribution and expression intensity of spatial marker genes on its different color channels. Initially, we applied a log1 transformation to gene expression representation. To validate the accuracy of our synthesized gene expression profiles, we specifically selected three spatial marker genes, displaying them on three distinct color channels, rather than using their low-dimension representation (e.g., the UMAP embedding). We used SOVIEW to clearly compare spatial discrepancies between the real and virtual slices, providing insights into their structural and expression variations.

